# MutaPhy: A clade-based framework to detect genotype-phenotype associations on phylogenetic trees

**DOI:** 10.64898/2026.04.19.719535

**Authors:** Amélie Ngo, Stéphane Guindon, Vincent Pedergnana

**Affiliations:** Virostyle - MIVEGEC (IRD, UM, CNRS), Montpellier, France; Méthodes et Algorithmes pour la Bioinformatique - LIRMM, Université de Montpellier, CNRS, Montpellier, France

**Keywords:** phylogenetics, genotype–phenotype association, GWAS, false positive rate, dengue virus, hepatitis C virus, evolutionary genomics

## Abstract

Understanding how genetic variation in pathogens influences clinical phenotypes observed in infected hosts is a fundamental challenge in evolutionary genomics and public health. Phenotypic traits such as infection severity are often non-randomly distributed within the pathogen’s phylogeny, suggesting the existence of evolutionary determinants but also violating the independence assumption underlying classical genome-wide association studies and potentially leading to inflated false positive rates. We present MutaPhy, a phylogeny-based method aimed at detecting correlations between a binary host phenotype and the corresponding pathogen genome by directly utilizing the hierarchical structure of phylogenetic trees. MutaPhy encompasses three different scales: (i) a subtree scale, on which relevant clades over-representing the phenotype of interest are detected using permutation-based tests; (ii) a tree scale, which agglomerates local signals into a global association statistics; and (iii) a site scale, whereby candidate mutational events on branches leading to significant clades are examined using ancestral sequence reconstruction. We evaluate the statistical behavior and detection performance of MutaPhy using simulations under diverse evolutionary scenarios. We also compare this tool to several existing phylogenetic association methods. As illustrative applications, we apply MutaPhy to dengue virus and hepatitis C virus datasets associated to clinical phenotypes in human hosts. Our results highlight the ability of the proposed approach to detect viral lineages associated to over-represented phenotypes while revealing limited evidence for robust mutation-level associations in these particular datasets. Altogether, MutaPhy provides a framework for guiding genotype–phenotype association analyses by leveraging phylogenetic structure, thereby reducing false positive findings and improving the interpretability of association signals.

## Introduction

A central goal in evolutionary biology and pathogen genomics is to determine how genetic variations in the pathogens’ genomes translates into clinical phenotypic variations observed in infected hosts. Recent large-scale epidemics, such as the COVID-19 pandemic, have illustrated how the successive emergence of viral variants can be associated with differences in transmissibility, immune escape or disease severity at the host level (Korber et al., 2020; Leung et al., 2021; Viana et al., 2022). In this context, identifying pathogen mutations associated with clinically relevant traits such as virulence, occurrence of specific symptoms, or resistance to treatment is essential for understanding disease mechanisms and improving surveillance strategies (Hadfield et al., 2018; Du Plessis et al., 2021).

Genome-wide association studies (GWAS) have become a standard framework for detecting genotype–phenotype associations in population genomic data. However, classical GWAS approaches rely on the implicit assumption of independence between sampled individuals. In practice, this assumption is often violated due to population stratification, where individuals share recent common ancestry, leading to correlations in genetic variation within cases and controls (Marchini et al., 2004; Astle and Balding, 2009; Price et al., 2010). Such structure can confound association signals and inflate false positive rates if not properly accounted for.

In evolutionary genomics of pathogens, the issue of population stratification is further exacerbated by the non-independence of the pathogen sequences themselves. Unlike classical population stratification, which is caused by shared ancestry within case and controls, pathogen genomes are related through a shared phylogenetic history, with mutations inherited along lineages. This induces strong correlations between samples, such that both genetic variants and associated phenotypes tend to cluster along specific branches of the phylogenetic tree (Felsenstein, 1985; Harvey and Pagel, 1991; Maddison, 1994; Earle et al., 2016; Saber and Shapiro, 2020). As a result, an association between pathogen genotype and host phenotype can often be explained by a single ancestral mutation inherited by multiple pathogen lineages, rather than by multiple independent mutation events. Ignoring this phylogenetic structure may therefore lead to misleading interpretations, where the same mutation appears to arise repeatedly across individuals, inflating false positive rates and complicating biological interpretation. (Cochrane et al., 2002; Marchini et al., 2004; Price et al., 2010; Earle et al., 2016).

Notwithstanding, the phylogenetic structure in pathogens’ genomes can be exploited to enhance association analyses. In a classical GWAS setting, associations are assessed by relating phenotype status (cases versus controls) and genetic state (mutant versus wild type). With the phylogeny, a third dimension can be incorporated by considering whether pathogen sequences belong to a clade enriched in the phenotype of interest or fall outside this clade. This additional clade-level information makes it possible to identify causal mutations in the pathogen’s genome, thereby providing a complementary approach to classical GWAS techniques that generally scour host genomes instead.

To address the limitations of classical GWAS in evolutionary settings, several methods have been specifically designed for pathogen genomic data. Among them, TreeWAS (Collins and Didelot, 2018) adapts GWAS principles to microbial populations by explicitly accounting for phylogenetic relatedness. TreeWAS relies on a binary representation of genetic variants and ancestral state reconstruction to test individual pathogen mutations for association with phenotypes defined at the pathogen level, while correcting for shared ancestry. However, this framework differs from our setting, where the objective is to identify associations between pathogen genetic variations and host clinical phenotypes. While effective at the mutation level, Tree-WAS focuses on individual variants and does not explicitly use the hierarchical organization of mutations along the phylogeny, which limits its ability to capture lineage-specific evolutionary signals.

Machine learning approaches such as random forests and neural networks have been explored to predict phenotypes, typically antibiotic resistance, from microbial genomic data (Libbrecht and Noble, 2015; Ren et al., 2022; Orcales et al., 2024). Although efficient, these methods typically lack interpretability and do not rely on an explicit evolutionary framework, making it difficult to relate predictions to the underlying evolutionary processes. Other techniques, which rely on phylogenies, have been developed to quantify non-random trait clustering within a tree. Metrics such as the Association Index (AI; Wang et al. 2001), Parsimony Score (PS; Fitch 1971) and Monophyletic Clade (MC; Salemi et al. 2005) statistics assess the degree of trait structuring on a tree, while BaTS (Parker et al. 2008) extends these approaches by accounting for phylogenetic uncertainty. More recently, CRP-Tree (Zhang et al. 2023) proposed a phylogeny-aware association test based on a Bayesian nonparametric model, designed to detect non-random clustering of a binary trait along a fixed phylogeny. While these methods are relevant for detecting global phylogenetic signal, they do not explicitly identify candidate mutations and do not provide a direct link between trait clustering and the evolutionary events that may underlie it. Moreover, they do not explicitly model how mutations along the phylogeny give rise to the observed distribution of traits.

In the present work, we introduce MutaPhy, a new clade-based framework for detecting geno-type–phenotype associations in a phylogenetic context. Instead of treating individual sequences as independent statistical units, MutaPhy considers evolutionary clades as the primary units of analysis. The general problem setting and the conceptual positioning of MutaPhy relative to existing phylogeny–trait association methods are illustrated in Figure 1. The method combines local tests of association across subtrees with ancestral sequence reconstruction to identify both phenotype clades and candidate mutations occurring on the branches leading to these clades. Our approach operates at three complementary scales: the subtree scale, where local association statistics and permutation *p*-values are computed; the tree scale, where signals are aggregated to assess global association; and the site scale, where mutations potentially contributing to the observed trait distribution are identified using ancestral sequence reconstruction. Importantly, MutaPhy uses the phylogenetic structure to focus on a smaller set of candidate mutations, instead of testing the whole genome. By restricting the analysis to these relevant sites, it reduces the number of tests performed, which helps increase statistical power and limit false positive results.

**Figure 1.**
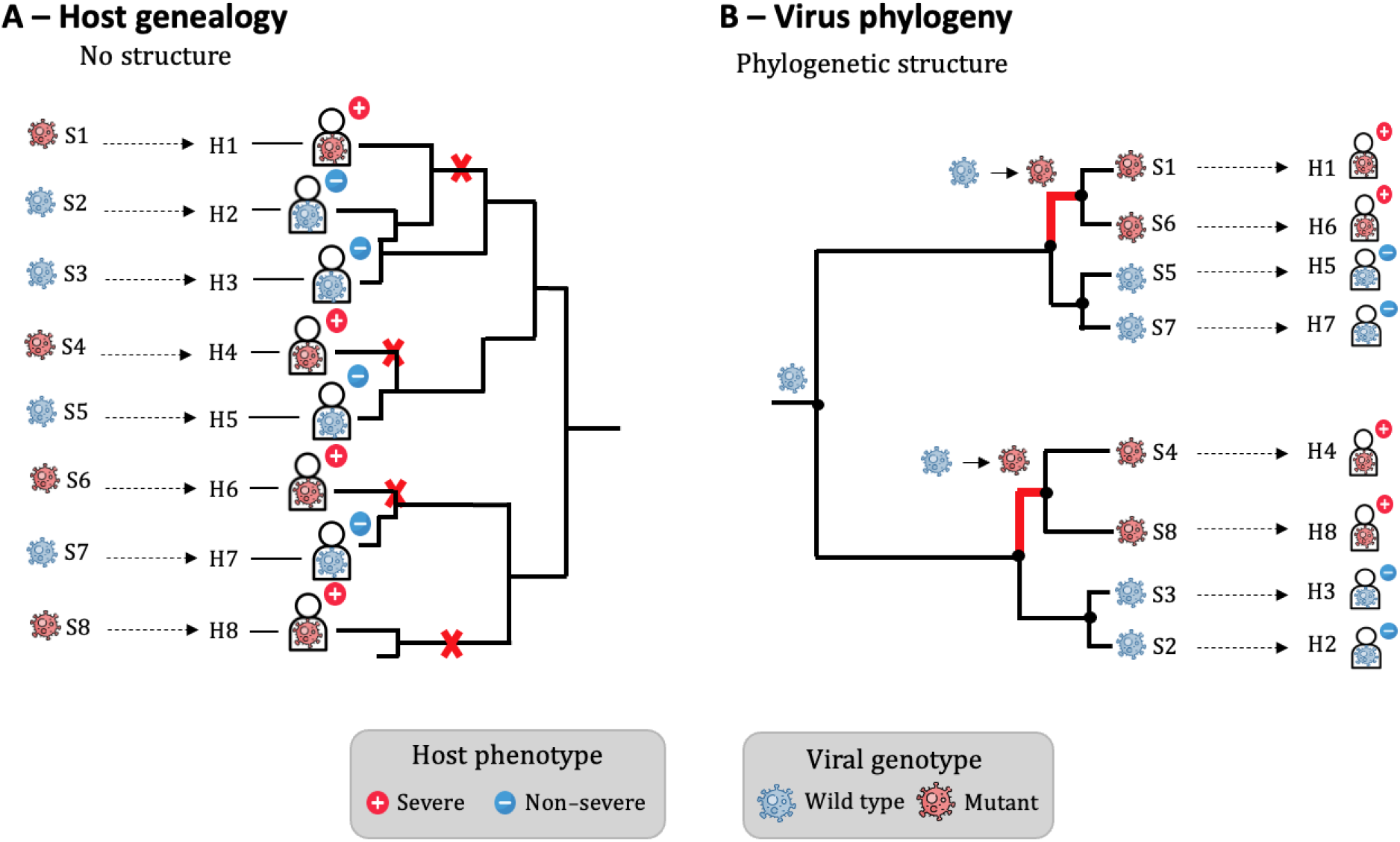
Illustration of genotype–phenotype associations in host and viral contexts. (A) Host genealogy showing no clear phylogenetic structure in the distribution of the phenotype: severe and non-severe cases are randomly distributed among related individuals. Explaining the observed association would require the same mutation to arise four times independently as indicated by the red crosses. (B) Viral phylogeny showing phylogenetic structure: pathogen sequences are related through a shared evolutionary history, and both mutations and associated host phenotypes tend to cluster along specific branches. In this case, only two mutation events are sufficient to explain the association, as highlighted by the red branches.

We evaluate the performance of MutaPhy using extensive simulations under a wide range of evolutionary scenarios and compare it to AI, PS, MC and CRP-Tree. Finally, we apply MutaPhy on real data from a clinical cohort of dengue virus infections, highlighting its potential for exploratory analysis of genotype-phenotype relationships in evolutionary genomics.

## Materials and Methods

### Overview

Figure 2 provides a schematic overview of the MutaPhy workflow and the articulation between its three scales of analysis. MutaPhy performs permutation of the phenotypes at the tips of the tree in order to test for an association at the subtree level. The corresponding *p*-values are corrected for non-independance between overlapping subtrees. These are agglomerated, yielding the statistics *p*_min_ and *p*_mean_ that characterize association at the tree level (see below), enabling comparison with existing tree-level methods (AI, PS, MC, CRP-Tree). In parallel, significant subtrees are used to identify candidate mutations at the level of individual sites via ancestral sequence reconstruction. A hypergeometric alternative test is also implemented as a complementary subtree-level association test.

**Figure 2.**
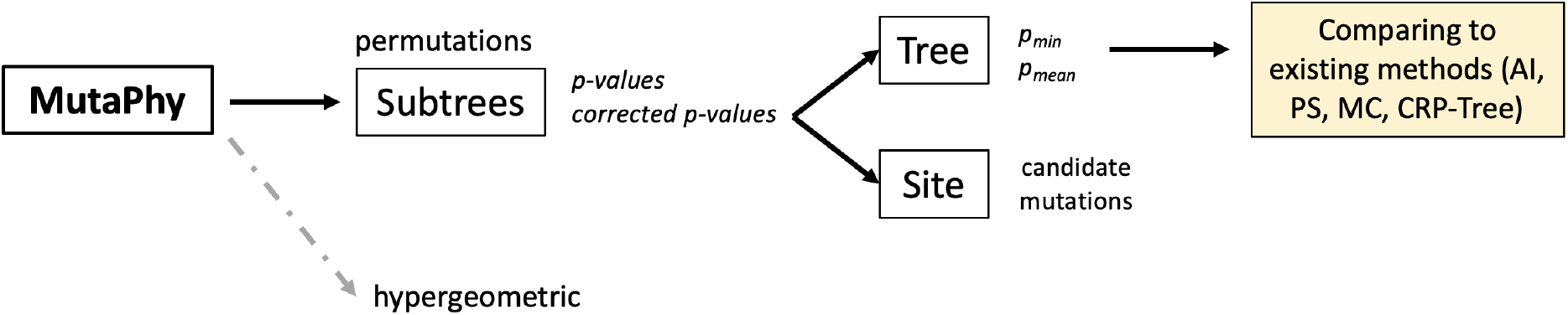
Overview of the MutaPhy workflow. See main text.

### Notation

Let *s* denote a sequence alignment. *s_ij_* is the character state (nucleotide or amino-acid) observed in the *i*-th sequence at the *j*-th position of the alignment matrix. We have *i* = 1, …, *n* and *j* = 1, …, *l* where *n* is the number of sequences in the sample and *l* is the length of the aligned sequences. *τ_n_* denote a rooted phylogenetic tree with *n* tips (and *n* − 1 internal nodes). Internal nodes are indexed by *u_i_, i* = *n* + 1, …, 2*n* − 1. Each internal node *u_i_* is the root of the subtree 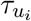 which contains 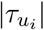 | tips. Each tip *u*_j_, *j* = 1, …, *n*, is associated with a binary phenotype *z_j_* ∈ {0, 1}. In the following, we will refer to *z_j_* = 1 as a “positive phenotype” and *z_j_* = 0 as a “null phenotype”. Let 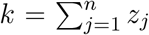 denote the number of tips with phenotype 1 in the whole tree, and 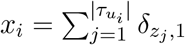 is the number of tips with phenotype 1 in subtree 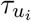.

### Subtree-level statistics

For each rooted subtree, MutaPhy tests whether the observed proportion of the positive (or null) phenotype deviates from that expected under the absence of phylogenetic structure. The null hypothesis states that any permutation of the phenotypes observed as the tips of the tree is equally probable, and that no mutation occurring on the branch(es) leading to the subtree being tested is associated with an increased probability of a particular phenotype. The alternative hypothesis states that the positive phenotype is over-(resp. under-)represented within the subtree of interest, suggesting that one or several mutations occurring on the branch(es) leading to this subtree may be associated with an increased probability of observing the 1 (resp. the 0) phenotype.

For each subtree 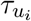, we thus evaluate the following statistic:

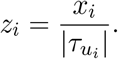

The core of MutaPhy relies on a permutation-based association test performed independently on each subtree. This test constitutes the primary statistical engine of the method. All subsequent steps, including tree-level aggregation and site-level mutation identification, are derived from the set of subtrees detected by this permutation procedure.

To test whether the positive phenotype is overrepresented in subtree 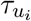, we generate an empirical null distribution by random permutation of the phenotypes across all tips in the tree. Specifically, let *N* be the number of these permutations and 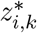 denotes the statistic for subtree 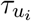 at permutation *k*. We use a one-sided test targeting an excess of positive phenotypes. The empirical *p*-value for subtree 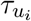 is:

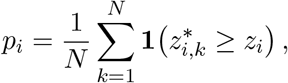

corresponding to the fraction of the simulations where the proportion of positive phenotypes in the subtree of interest is greater or equal to the observed in the original data set. For computational convenience, we define a binary matrix *B* ∈ {0, 1}^*N*×(*n*−1)^, where rows correspond to permutations and columns correspond to all subtrees and

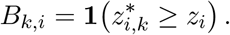

Figure 3 provides a schematic illustration of this construction, showing how permutation outcomes are mapped to the binary matrix *B* and how the corresponding 0/1 values can be visualized directly on the phylogenetic tree for subtrees.

**Figure 3.**
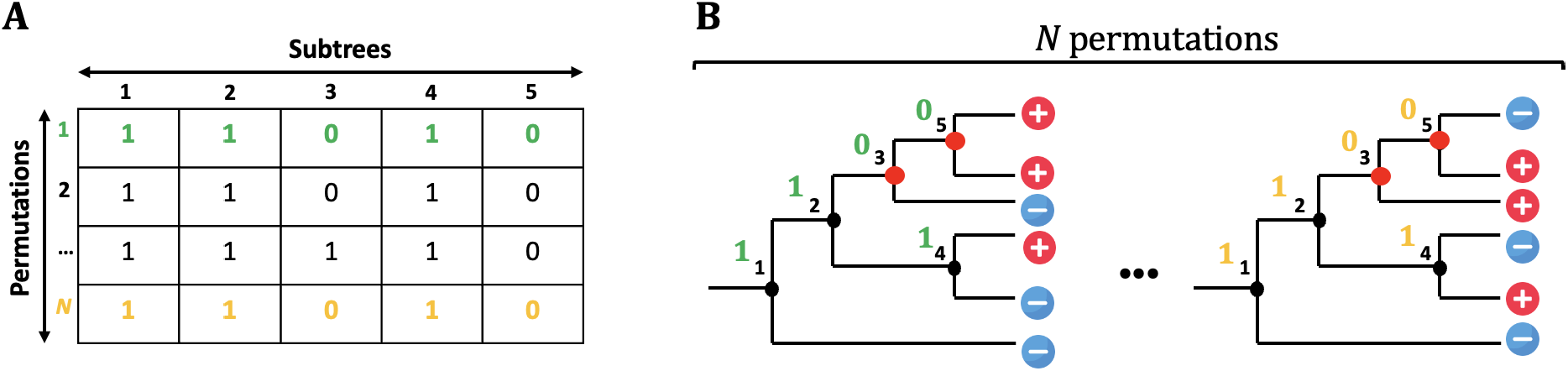
Illustration of the binary permutation matrix construction. (A) Example of a binary matrix *B* obtained from *N* permutations, where each column corresponds to a rooted subtree and each row to a permutation. Only a subset of permutations is displayed for visualization purposes. (B) Visualization of the same permutation outcomes directly on the phylogenetic tree. Colored disks for each internal node indicates whether the corresponding rooted subtree is declared significant (1, red) or not (0, black) after permutation testing.

The *p*-value of a given subtree may also be obtained more directly by considering sampling without replacement 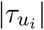 balls in an urn containing *m* balls of positive type and *n* − *m* balls of null type. The probability of observing *x* balls of positive type is then simply:

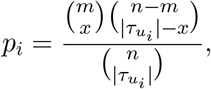

corresponding to an hypergeometric distribution. While this parametric approach is valid (and computationally efficient), its application over the entire set of subtrees assumes their independence, an assumption that is violated in phylogenetic settings due to the nested and overlapping structure of subtrees. This limitation motivates the use of the permutation test in MutaPhy, which naturally accounts for these dependencies.

In order to avoid redundant detections and explicitly accommodate for phylogenetic structure, MutaPhy applies an iterative filtering procedure among subtrees that satisfy *p_i_ < α*. Significant subtrees are ranked by increasing *p_i_* values. Ties are resolved by giving priority to larger subtrees over smaller ones. At each iteration, the top-ranked subtree is selected, and all remaining significant subtrees that are either ancestors or descendants of the selected subtree are considered redundant. For those redundant subtrees, all entries equal to 0 in the corresponding columns of the binary matrix *B* are replaced by 1, which increases their empirical *p*-values, effectively removing them from the final set. This procedure is repeated until no overlapping (ancestor/descendant) significant subtrees remains.

This redundancy-correction mechanism is illustrated in Figure 4. In this example, two nested subtrees (rooted at nodes 3 and 5) are initially detected as significant. Because subtree 5 is fully contained within subtree 3, it does not represent an independent evolutionary signal. The filtering procedure therefore modifies the corresponding column of *B* by replacing all remaining 0 entries with 1, thereby inflating its permutation *p*-value above the significance threshold.

**Figure 4.**
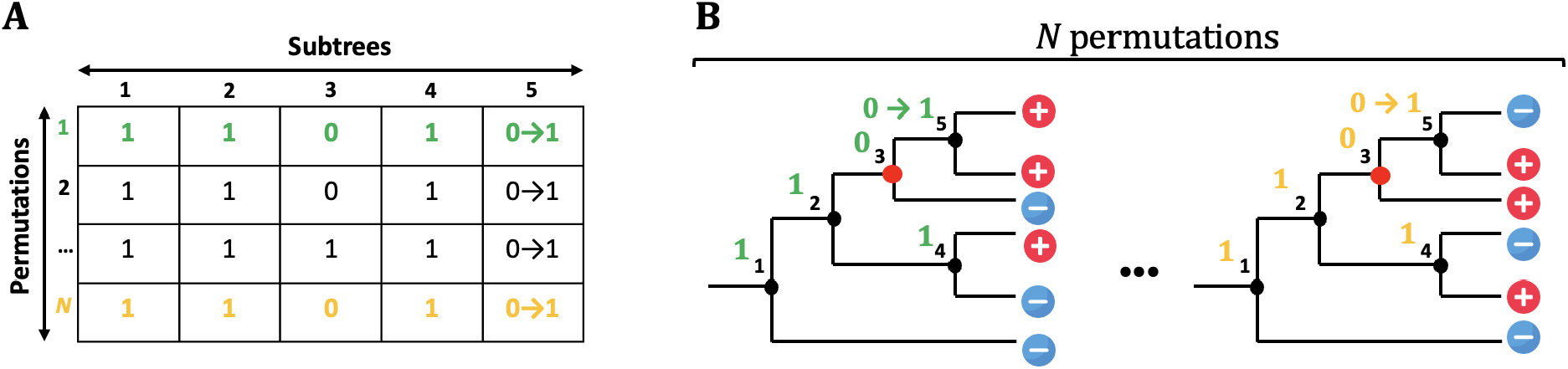
Illustration of the binary permutation matrix *B* and redundancy filtering. (A) Green and orange arrows illustrate the redundancy correction step for permutation 1 and N, where 0 → 1 replacements are applied to nested subtrees. (B) Subtrees rooted at nodes 3 and 5 are detected as significant (red), but because subtree 5 is nested within subtree 3, it is considered redundant and corrected by the filtering procedure.

Moreover, testing all subtrees induces multiple comparisons. A classical Bonferroni correction using the total number of tested subtrees would be overly conservative because subtree tests are not independent. We therefore use a modified Bonferroni correction based on the maximum number of independent signals that can be detected, denoted as *m* and defined as the maximal number of non-overlapping significant subtrees returned by the redundancy-filtering procedure. We then define an adjusted threshold

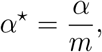

and retain subtrees whose corrected *p*-values remain below *α*^⋆^ after the redundancy correction applied to *B*. After redundancy filtering and multiple-testing correction, a subtree 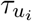 is declared *positively associated* with the phenotype if its corrected *p*-value satisfies

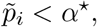

The set of positively associated subtrees constitutes the input for both tree-level aggregation and site-level mutation identification.

### Tree-level statistics

To summarize phenotype–genotype association at the whole-tree level, MutaPhy relies on two summary statistics derived from subtree-level permutation tests. The minimum subtree statistic is defined as

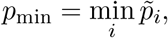

and captures the presence of a strong and localized association signal. The second statistic, the mean subtree statistic, is defined as

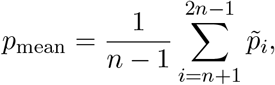

where *n* − 1 is the number of internal nodes. This statistic captures more diffuse association patterns across the phylogeny. These statistics are used as tree-level summary scores to compare MutaPhy with alternative phylogenetic association methods (AI, MC, PS and CRP-Tree) in simulation studies.

### Identification of causal mutations

For each significant subtree, we identify candidate mutations on the branch leading to the root of the corresponding clade. Ancestral nucleotides are reconstructed at internal nodes for each site of the alignment using maximum likelihood (function `ace` in the `ape` R package (Paradis and Schliep, 2019)). For each site, we compare the reconstructed state at the root of the clade and its direct parent. A mutation is recorded when the most probable nucleotide differs between these two nodes.

In order to handle ambiguous reconstructions, we distinguish between clear state changes and uncertain cases. A mutation is recorded as a state change if either the most probable nucleotide differs between the root of the clade and its direct ancestor, or if the probability of the most probable state at the root differs from that observed in its direct ancestor by 0.3 or more. In all other cases, the mutation is considered as ambiguous, reflecting uncertainty about whether a mutation occurred on the branch leading to the clade. This threshold of 0.3 was chosen empirically to balance sensitivity and specificity in simulated data. While ancestral state reconstruction provides a practical way to identify candidate mutations, a more rigorous approach would rely on mutation mapping methods that explicitly model substitution events along branches (Nielsen, 2002). However, these approaches are computationally demanding, hence our decision to opt for a simpler solution.

When multiple non-overlapping clades are detected, the site-level output is defined as the union of the sets of candidate mutations identified on the branches leading to each clade. The intersection of the sets of candidate mutations across independent clades may further reveal convergent mutations: identical nucleotides (or amino-acids) that have arisen independently in distinct lineages and are associated with the same phenotype.

### Simulation design and evaluation

#### Objectives

We evaluated MutaPhy using simulations in which phylogenetic trees, evolving nucleotide sequences, and host phenotypes were generated synthetically. In simulated data, the causal mutations underlying the phenotype are known by construction, which enables a direct assessment of (i) the detection of association at the tree level, (ii) the identification of associated subtrees, and (iii) the recovery of causal mutations on branches leading to clades detected in (ii).

#### Phylogenetic trees and sequence evolution

Phylogenetic trees were simulated under the Kingman coalescent (Kingman, 1982) using rcoal() from the ape R package (Paradis and Schliep, 2019). Starting from an ancestral sequence at the root of each tree, nucleotide evolution evolved under the Kimura two-parameter substitution model (K80) (Kimura, 1980), with transition/transversion ratio set to 4. Branch lengths were used as relative evolutionary times, and the number of substitutions occurring on a branch of length *l* was drawn from a Poisson distribution with parameter *λ* = *µ* × *l*, where *µ* = 10^−4^ is the per site mutation rate, chosen to be consistent with typical RNA virus mutation rates. In our setting, *µ* primarily acts as a scaling factor for the total number of substitutions. To ensure comparable signal intensity across scenarios, genomes were grown progressively by adding sites until a fixed number of mutations was reached (*N*_mut_ = 100 per tree), so that all simulated datasets contained the same total number of mutational events regardless of the final genome length.

Under the null hypothesis, mutations do not determine the phenotype. Under the alternative hypothesis, a controlled fraction of simulated mutations, *M*_tot_ ∈ [0, 1], was designated as causal mutations. For each simulated dataset, the *N*_mut_ mutations were ranked according to their position in the alignment, and the first ⌊*M*_tot_ × *N*_mut_⌋ mutations were designated as causal, while the remaining mutations were considered non-causal. For each tip *i*, we computed an infectivity score *s_i_* defined as the number of causal mutations accumulated along the path from the root to *i*. This score was mapped to a probability of observing phenotype 1 via a logistic function:

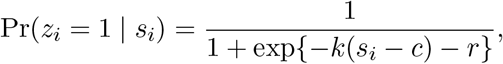

with *c* controlling the inflection point and *k* the slope. In order to model background (non-genetic) variability, we enforced a baseline probability Pr(*z_i_* = 1 | *s_i_* = 0) = *b* through the offset term *r*, considering both low-noise (*b* = 0.01) and noisy (*b* = 0.20) settings. To explicitly control background (non-genetic) variability, we constrained the logistic model to enforce a predefined baseline probability *b* of phenotype expression in the absence of causal mutations, i.e. *P* (*z_i_* = 1 | *s_i_* = 0) = *b*. This was achieved by introducing a bias term *r* determined analytically so that the constraint is satisfied (see Supplementary Method S1). Finally, phenotypes were sampled independently at the tips as *Z_i_* ∼ Bernoulli(*p*). The effect of the causal mutation proportion *M*_tot_ and of the background noise parameter *b* on the phenotype–mutation mapping is illustrated in Supplementary Figure S1.

We varied the number of tips (*n* ∈ {20, 50, 100, 300}), the proportion of causal mutations (*M*_tot_ ∈ {1%, 10%, 50%}), the presence or absence of background noise (*b* ∈ {0.01, 0.2}), and the hypothesis (*H*_0_ vs. *H*_1_). For each parameter combination, 100 trees were simulated, yielding 48 scenarios in total. All simulation parameters are summarized in Supplementary Table 1. We compared MutaPhy to four widely used phylogeny-trait association methods: Parsimony Score (PS) (Fitch, 1971), Association Index (AI) (Wang et al., 2001), Monophyletic Clade (MC) (Salemi et al., 2005), and CRP-Tree (Zhang et al., 2023). All methods were applied to the same simulated phylogenetic trees *τ_n_* and phenotype vectors *Z*.

The Parsimony Score (Fitch, 1971) measures the minimum number of phenotype state changes required to explain the observed distribution of phenotypes at the tips of the tree. Significance is assessed by comparing the observed score to an empirical null distribution obtained by random permutations of tip phenotypes. Lower scores indicate stronger phylogenetic clustering than expected by chance if no phylogenetic structure is present.

The Association Index (Wang et al., 2001) quantifies the degree of phenotype clustering across all internal nodes of the tree. For each internal node *u_i_*, let *f_i_* denote the number of tips in 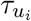 sharing the most frequent phenotype. The AI statistic is defined as

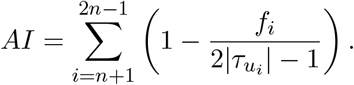

Lower *AI* values correspond to stronger phylogenetic structure. Statistical significance is assessed by comparing the observed *AI* to its empirical null distribution obtained by phenotype permutations.

The Monophyletic Clade (Salemi et al., 2005) statistic evaluates whether individuals sharing the same phenotype form large monophyletic groups (Wang et al., 2001). Let

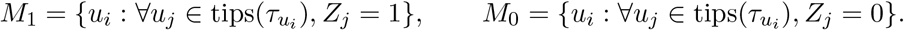

The MC statistics are defined as

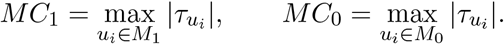

Significance is evaluated by comparing observed *MC*_1_ and *MC*_0_ values to permutation-based null distributions.

Finally, CRP-Tree (Zhang et al., 2023) is a phylogenetic association test for binary traits based on a generative model inspired by the Chinese Restaurant Process (CRP). Unlike previous methods (PS, AI, MC), CRP-Tree relies on a probabilistic framework that jointly considers a fixed phylogenetic topology *τ_n_* and an ordered insertion process compatible with this topology. The method does not generate new phylogenies but instead explores multiple planar representations of the observed tree, corresponding to different hypothetical orders in which tips could have been inserted during evolution. Given a phenotype vector *Z* = (*Z*_1_, …, *Z_n_*), tips are considered sequentially according to a planar ordering. At each insertion step *k*, let *w_k_* denote the number of previously inserted tips sharing the same phenotype as *Z_k_*. Under the CRP-inspired model, the theoretical probability of preferential attachment to a tip of the same phenotype is given by

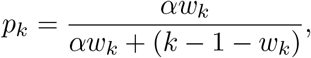

where *α* ≥ 1 controls the strength of phenotype clustering, with *α* = 1 corresponding to a neutral model without association.
For a given planar representation, CRP-Tree defines a clustering statistic *S* as the number of insertion steps for which attachment to a tip of the same phenotype is possible (i.e., *w_k_ >* 0). The observed statistic *µ*_0_ is obtained by averaging *S* over multiple random planar representations of the observed tree for a fixed *α >* 1. Statistical significance is assessed by comparing *µ_0_* to an empirical null distribution generated from random permutations of phenotype labels under the neutral model (*α* = 1).

### Evaluation criteria and summary statistics

For MutaPhy, tree-level evidence was summarized from subtree-scale permutation *p*-values using *p*_min_ and *p*_mean_, which served as global scores to discriminate tree generated under the alternative vs. the null hypotheses. Performance was assessed using ROC curves and AUC.

At the subtree level, we evaluated MutaPhy’s ability to identify phenotype-associated clades by comparing detected significant subtrees to the subtrees corresponding to branches where causal mutations were introduced in the simulations. Because subtree positives are typically rare, performance was summarized using precision-recall curves and AUC-PR. We additionally reported results under two evaluation definitions: a strict definition, where a detection is counted as correct only if the causal mutation occurs exactly on the branch leading to the detected sub-tree, and a hierarchical definition, where descendants of a causal clade are also considered correct in order to account for inheritance along the tree. At the site level, we evaluated the identification of candidate mutations by comparing mutations returned by ancestral reconstruction on branches leading to detected clades to the set of truly causal mutations, using precision and recall. Site-level evaluation was restricted to simulations in which at least one significant subtree was detected, since no candidate mutations can be produced otherwise.

## Results

### Tree-level detection

We first assessed the ability of MutaPhy to detect an association between host phenotype and pathogen phylogeny at the whole-tree level using simulations. For each simulated tree, permutations of phenotype labels at the tips were summarized using the two statistics *p*_min_ and *p*_mean_, where *p*_min_ is the smallest *p*-value across all subtrees and *p*_mean_ is the average *p*-value (see Methods). These statistics were used to discriminate trees simulated under the null hypothesis of no association from those simulated under the alternative hypothesis and compared to AI, PS, MC and CRP-Tree using ROC curves and AUC.

Using the *p*_min_ statistic (Figure 5), MutaPhy achieves competitive performance under noiseless conditions (AUC = 0.835), comparable to AI (0.819) and PS (0.893), but slightly below CRP-Tree (0.909). However, in the presence of background noise, its performance drops markedly (AUC = 0.584), indicating a strong sensitivity when relying on the most extreme subtree *p*-value. In some regions of the ROC curve, MutaPhy with *p*_min_ falls below the random baseline. This occurs because *p*-values falling below a nominal threshold under the null hypothesis are likely to arise by chance when considering multiple tests, even in situations where the null hypothesis applies. A similar behavior is also observed for CRP-Tree, suggesting that both methods may produce unstable rankings under noisy conditions, in contrast to PS, AI and MC, whose global statistics remain more stable. In contrast, when the tree-level statistic is summarized using *p*_mean_ (Figure 6), MutaPhy becomes the best-performing method in noiseless conditions (AUC = 0.944), outperforming all competing approaches. Importantly, it is also more robust under noisy conditions (AUC = 0.752), ranking among the top-performing methods and clearly improving over its *p*_min_ formulation. Altogether, these results show that aggregating subtree-level evidence through *p*_mean_ provides a more stable and robust global test, whereas reliance on extreme subtree signals (*p*_min_) leads to reduced specificity in noisy settings. From an applied perspective however, *p*_min_ remains suitable as it directly highlights the most strongly associated subtree and better reflects the method’s primary objective of pinpointing clades of interest in the pathogen phylogeny.

**Figure 5.**
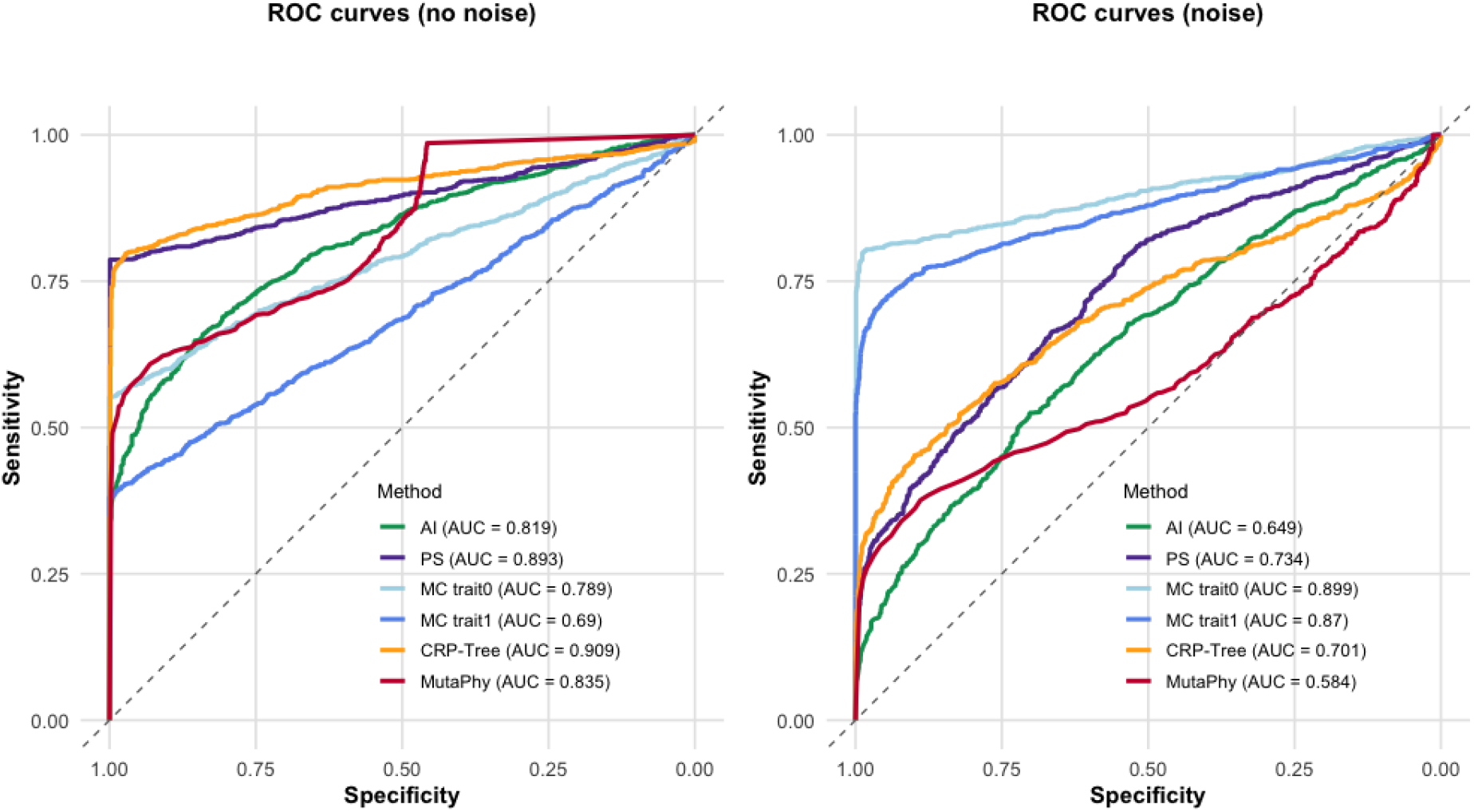
ROC curves comparing MutaPhy using the *p*_min_ summary statistic to existing methods (AI, PS, MC, and CRP-Tree), under noiseless (left) and noisy (right) simulation scenarios

**Figure 6.**
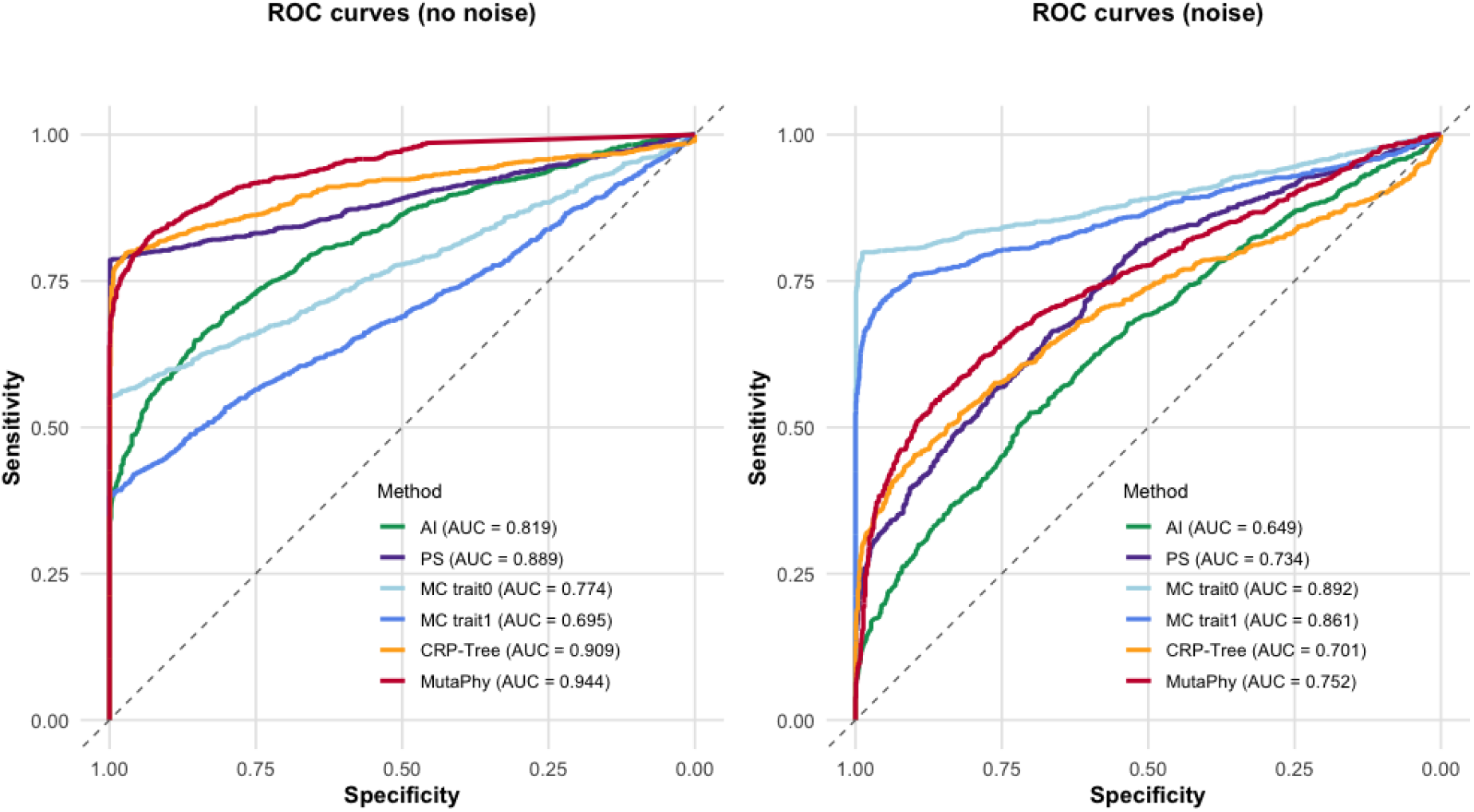
ROC curves comparing MutaPhy using the *p*_mean_ summary statistic to existing methods (AI, PS, MC, and CRP-Tree), under noiseless (left) and noisy (right) simulation scenarios

### Subtree-level identification

We next evaluated the ability of MutaPhy to identify clades with over-represented positive phenotypes. For each simulation scenario, 100 independent trees were generated, and permutation *p*-values were computed for all rooted subtrees. Because phenotype-associated clades are rare relative to the total number of tested subtrees, the accuracy of our approach was assessed using precision–recall (PR) curves and the area under the PR curve (AUC-PR). By default, results are reported using *p*-values deriving from permutations and corrected for non-independence plus multiple testing (raw results are provided in Supplementary Figure S2).

Figure 7 summarizes performance under both the strict (i.e., *p*-values are not corrected using the tree structure) and the hierarchical (*p*-values are corrected) definitions. Under the null hypothesis, specificity was consistently close to 1 across all simulation settings, indicating a low false positive rate. Under the alternative hypothesis, the hierarchical approach consistently outperformed the strict one. Near-perfect performance was achieved for moderate to high proportions of causal mutations (*M*_tot_ ≥ 10%) in noiseless settings. Multiple-testing correction slightly reduced sensitivity under the strict approach and for very low causal proportions, but did not substantially affect the hierarchical method. Overall, these results indicate that MutaPhy performs well in identifying clades where a given phenotype is over-represented, with the hierarchical approach providing the most stable performance across tree sizes and noise levels.

**Figure 7.**
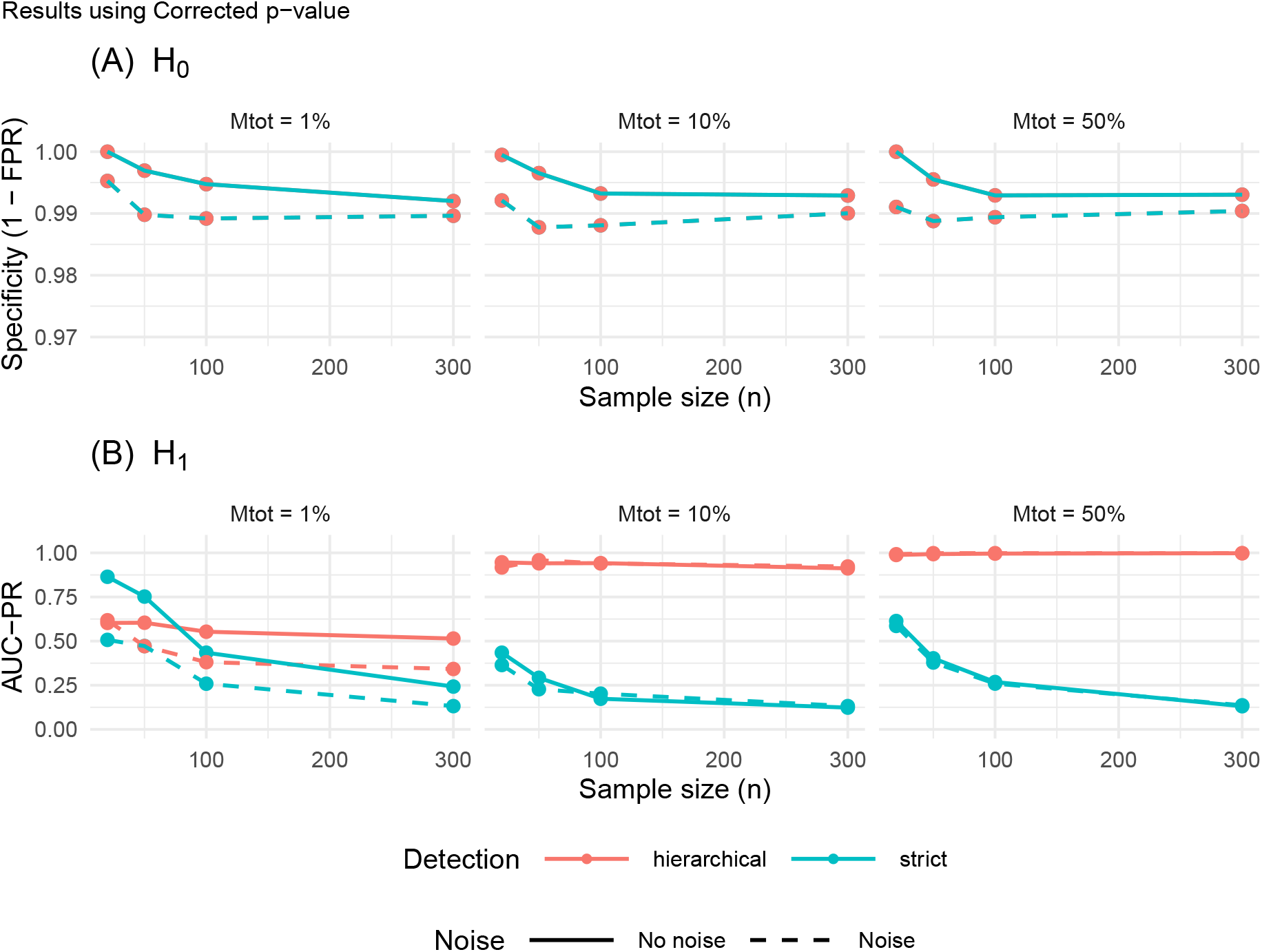
Subtree-level performance of MutaPhy using multiple-testing–corrected permutation *p*-values, under strict and hierarchical evaluation definitions.

We additionally evaluated a hypergeometric subtree test as a complementary validation of the non-parametric permutation-based approach. Across simulation settings, both approaches show comparable performance, with specificity consistently close to 1 under the null hypothesis (typically above 0.98 for both methods) and similar AUC-PR values under the alternative hypothesis, with only minor differences observed across scenarios. For low signal scenarios (*M*_tot_ = 1%), the hypergeometric test shows slightly higher AUC-PR values, likely due to its more permissive behavior. Results are reported in Supplementary Figure S3. However, the permutation-based approach provides important methodological advantages. In contrast to the hypergeometric test, it takes into account the phylogenetic dependence between sequences and can handle overlapping subtrees through the binary matrix update. Besides, this allows a proper multiple-testing correction, which is not possible with the hypergeometric test.

### Site-level identification of candidate mutations

MutaPhy can be used to identify candidate mutations associated to the phenotype of interest. To this end, we adopt a simple strategy: for each significant subtree, all mutations occurring on the branch leading to the detected clade are considered as potential candidates. This rationale is inherently naive as it does not attempt to distinguish causal mutations from neutral ones. Rather, it provides an exhaustive list of mutations associated with the emergence of the phenotype-enriched clade. The proposed approach is thereby mainly geared towards exploratory analysis rather than pinpointing causal mutations with high precision (i.e., with a low false positive rate).

Because the identification of candidate mutations requires that at least one significant sub-tree is detected, site-level evaluation was restricted to trees for which MutaPhy detected at least one phenotype-associated subtree. Across simulation scenarios, precision increases with the proportion of causal mutations (*M*_tot_), but remains moderate overall. When causal mutations are rare (*M*_tot_ = 1%), precision is very low (typically below 0.1), reflecting a high proportion of non-causal candidates. For intermediate scenarios (*M*_tot_ = 10%), precision increases to approximately 0.1–0.3 depending on sample size and noise conditions. In contrast, when a large fraction of mutations are causal (*M*_tot_ = 50%), precision reaches values around 0.5–0.7, indicating that a substantial proportion of detected mutations are truly associated with the phenotype (Supplementary Figure S4). Despite this moderate precision, MutaPhy considerably reduces the search space, restricting downstream analyses to a limited number of candidate mutations instead of the full set of genomic sites. In terms of recall, performance depends on how the causal signal is distributed across the phylogeny. When causal mutations are rare (*M*_tot_ = 1%), recall is highly variable across replicates, reflecting whether the single causal mutation falls within a detected clade. As the number of causal mutations increases (*M*_tot_ ≥ 10%), recall decreases overall, particularly in scenarios with larger sample sizes or noise. This reflects the increasing difficulty of recovering all causal mutations when they are distributed across multiple independent branches of the tree (Supplementary Figure S5).

To further evaluate the relevance of our approach, we compared the set of causal mutations detected with MutaPhy to that obtained with genome-wide association tests. More precisely, we performed standard association tests (Fisher’s exact test or chi-square test) at all genomic positions and compared the set of significant sites to that derived from MutaPhy (Figure 8). As expected, the number of false positives detected by global GWAS increases with the number of sequences and is particularly high when causal mutations are rare (*M*_tot_ = 1%). This is due to the large number of tests performed across the genome. In contrast, MutaPhy strongly reduces the number of false positives across all simulation scenarios, with a reduction of up to a factor of 3 in some settings (*n* = 300). This reduction is accompanied by a decrease in the number of true positives, but the loss remains limited compared to the reduction in false positives, resulting in a net gain in precision. As a result, precision is slightly improved when using MutaPhy. In addition, the number of false positives varies strongly with sample size for the global GWAS, while it remains more stable with MutaPhy. These results are confirmed when considering *n* = 300, where MutaPhy reduces false positives by about 63% and increases precision by approximately 3.5 points, while still recovering around 46% of the true positives identified by the global GWAS.

**Figure 8.**
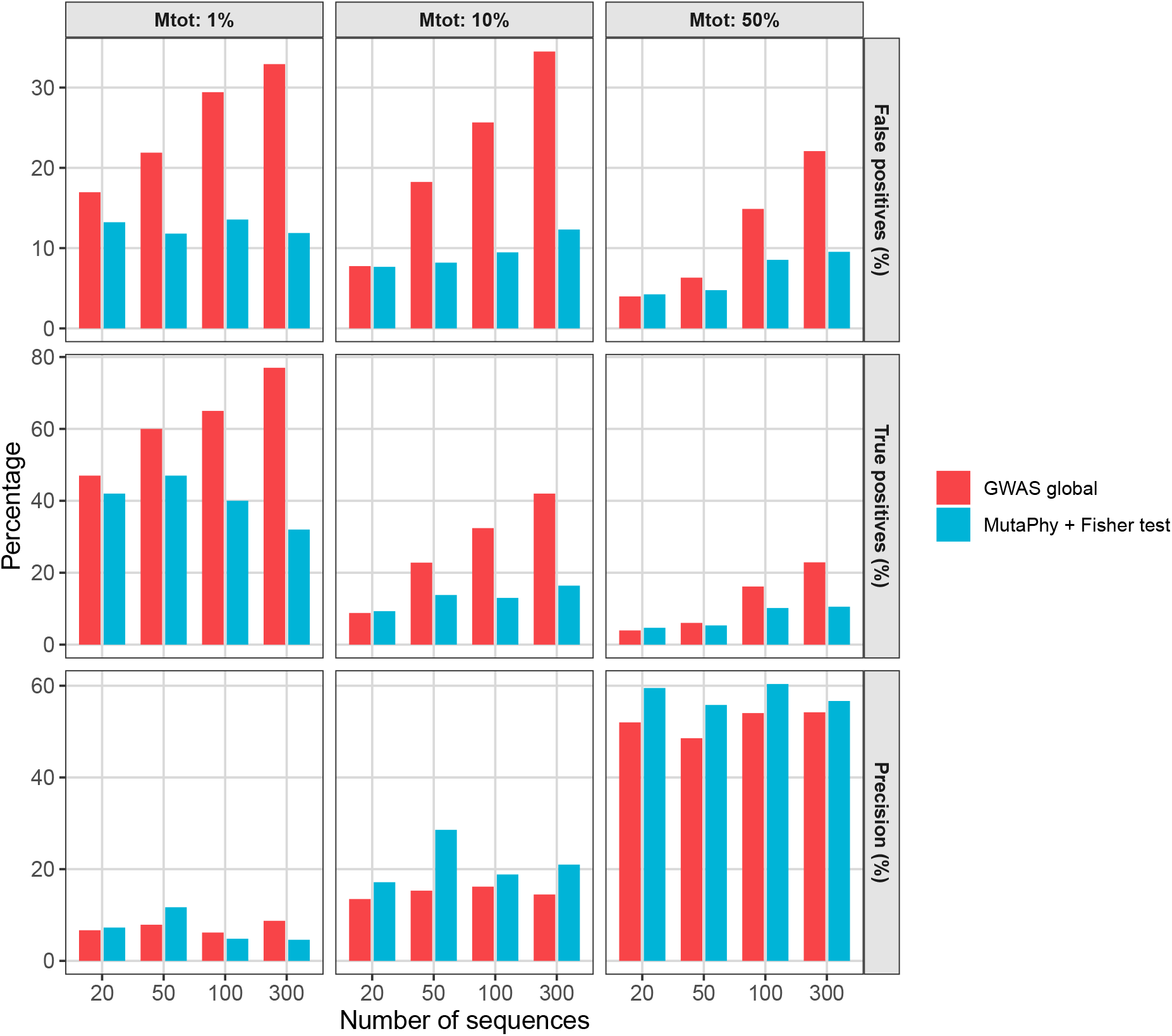
Comparison of a genome-wide association strategy (GWAS global) and MutaPhy with targeted association testing across simulation scenarios. Results are shown for different proportions of causal mutations (*M*_tot_) and number of sequences. For each setting, the percentage of false positives, true positives, and precision are reported. Results are shown for noiseless simulations; similar trends were observed under noisy conditions.

### Application to hepatitis C virus clinical data

To test the performance of MutaPhy in real conditions, we applied it to a dataset of 507 hepatitis C virus (HCV) genomes (genotype 3a), with treatment outcome defined as a binary phenotype (91 non-SVR and 416 SVR patients), where SVR (sustained virological response) corresponds to the absence of detectable virus after antiviral treatment, as described in Smith et al. (2021). In this study, the author performed a viral genome-wide association study of HCV sequences and identified three viral polymorphisms significantly associated with SVR. We inferred phylogenetic relationships using IQ-TREE2 (Minh et al., 2020). At the subtree level, MutaPhy identified several phenotype-enriched clades. When testing for clades enriched in SVR, seven significant clades were detected, including six small clades of size two and one clade of size four. Aggregating mutations located on the branches leading to these clades resulted in 429 candidate mutations. Among these, only four mutations were associated with the phenotype at a false discovery rate (FDR) of 15%, and each was supported by a single clade of size two. The three amino acid variants (NS2: alanine at position 119 and valine at position 132 and NS3: valine at position 67) previously reported (Smith et al., 2021) to be associated with SVR outcome were not detected by MutaPhy. These variants seem to appear several times independently across the tree, rather than being grouped within clades large enough to be detected by MutaPhy, making them difficult to detect with our clade-based approach (Figure 9). In addition, it is likely that mutations favoring non-SVR occurred during the course of treatment within patients and thus could not be identify by ancestral reconstruction. Overall, the limited number of robust associations likely reflects both reduced statistical power, due to the small size of detected clades, and the multifactorial nature of treatment response in HCV infection. Indeed, sustained virological response is influenced by a combination of viral and host factors, including host genetics, immune response, and treatment-related variables, which may hide direct associations with individual viral mutations as observed in dengue analysis.

**Figure 9.**
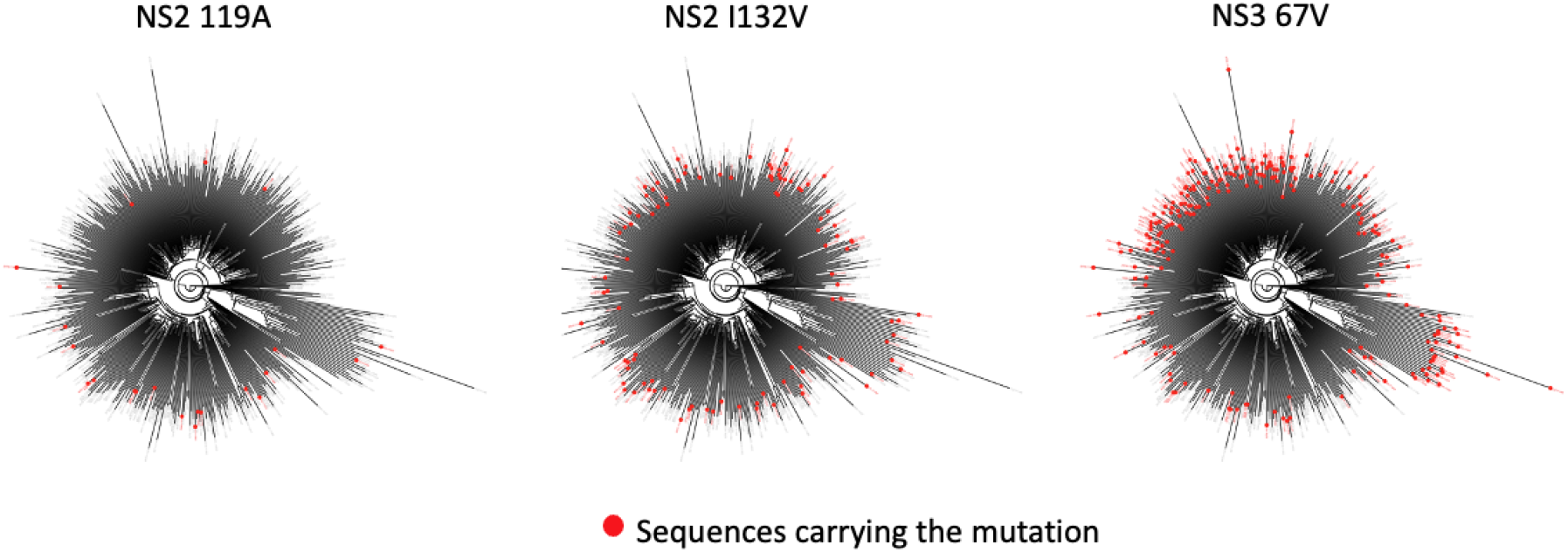
Phylogenetic distribution of three amino acid variants previously reported (Smith et al., 2021) to be associated with treatment outcome (NS2 119A, NS2 I132V, and NS3 67V)

## Discussion

A central challenge in pathogen GWAS is controlling false positives caused by the non-independence of pathogen sequences. Because they share a common evolutionary history, pathogen mutations and the associated host phenotype tend to cluster along the phylogeny. Ignoring this structure can lead to spurious associations, where mutations appear linked to the host phenotype simply due to shared ancestry rather than true causal effects.

In this study, we introduce MutaPhy, a clade-based framework designed to detect genotype-phenotype associations in phylogenetic settings while explicitly accounting for shared ancestry. Unlike existing tree-level association statistics (AI, PS, MC, CRP-Tree), MutaPhy identifies phenotype clades and corresponding candidate mutations through tip-label permutation of host phenotypes together with ancestral pathogen sequences reconstruction. By restricting the analysis to evolutionarily relevant regions, MutaPhy reduces the number of tests and guides down-stream association analyses in a more targeted and interpretable way, thereby limiting false positives.

Although *p*_mean_ achieves higher AUC values at the tree level, the *p*_min_ statistic is more closely aligned with the main goal of MutaPhy. By focusing on the single most strongly associated subtree, *p*_min_ directly points to identified phylogenetic regions where the phenotype of interest is overrepresented, which is central to the clade-based design of the method. In practice, this implies a trade-off between global discrimination and interpretability. While *p*_mean_ is well suited to assess whether an overall association is present across the tree, *p*_min_ is more appropriate for identifying specific clades that may carry a biologically meaningful signal. Rather than representing competing choices, these two summaries play complementary roles within MutaPhy.

At the site scale, MutaPhy successfully recovers isolated causal mutations but shows decreasing recall when multiple causal events are distributed across different branches (as highlighted by our application of the method in HCV data). This behavior highlights a fundamental limitation of identifying all causal variants in polygenic evolutionary architectures. Importantly, MutaPhy is designed as an exploratory tool: even with moderate precision, it drastically reduces the search space from thousands of sites to a manageable set of candidate mutations, enabling focused biological validation rather than definitive causal inference.

We showed that combining MutaPhy with classical association tests substantially improves the balance between sensitivity and specificity. In scenarios with large sample sizes (*n* = 300), our combined approach reduces false positive detections by up to 63%, while retaining approximately 46% of the true positives identified by a standard GWAS. Importantly, this trade-off results in an overall gain in precision (+3.5%), indicating that the reduction in spurious associations more than compensates for the loss in sensitivity. These results support the use of MutaPhy as a filtering step prior to association testing, enabling a more targeted and interpretable analysis of genotype-phenotype relationships.

Applications to a real dataset further illustrate both the potential and the limitations of the approach. Analyses of hepatitis C virus data did not reveal robust mutation-level associations after statistical correction. While these negative results may appear disappointing, they are consistent with the biological complexity of the phenotype. In particular, HCV treatment response is strongly influenced by host-related factors, including immune history and environmental context, which are not captured by viral genomic variation alone. These findings highlight an important point: the absence of significant mutation-level associations does not invalidate the phylogenetic signal detected at the clade level, but rather suggests that the underlying mechanisms may be multifactorial or involve weak genetic effects distributed across the genome.

Several limitations of MutaPhy should be acknowledged. First, statistical power decreases for small clades or rare phenotypes, as detection relies on identifying significant clustering within subtrees. Second, phylogenetic uncertainty is not explicitly modeled, as the tree is assumed to be fixed. Third, subtree-level tests are not independent due to the nested structure of phylogenies, and although hierarchical filtering reduces redundancy, formal multiple-testing correction under dependency remains an open challenge. Finally, candidate mutation identification is restricted to branches leading to the most significant clades, which favors interpretability but may miss deeper or more diffuse evolutionary signals. Future developments could address these limitations by incorporating host-related covariates, extending the framework to continuous or multi-class traits, and integrating recombination-aware models. More broadly, combining phylogeny-aware filtering with association testing represents a promising direction to improve the robustness and interpretability of genotype-phenotype analyses in pathogen genomics.

In summary, MutaPhy is a clade-based framework for phylogeny–phenotype association testing that links localized phenotype clustering to candidate mutational events on the branches defining associated lineages. Through extensive simulations, we show that MutaPhy robustly identifies phenotype-overrepresented clades and provides an interpretable reduction of the candidate search space at the site level. When combined with standard association tests, this approach substantially reduces false positives while maintaining a meaningful fraction of true signals, leading to an overall improvement in precision. Applications to HCV dataset highlights both the potential and the limitations of the framework: while phylogenetic structure can reveal lineage-level signals, mutation-level associations remain difficult to detect for complex, multifactorial phenotypes. Overall, MutaPhy provides an exploratory strategy to guide association testing by prioritizing evolutionarily relevant regions of the genome, thereby improving the robustness and interpretability of genotype–phenotype analyses in pathogen genomics.

## Code and data availability

The MutaPhy implementation, together with the simulation framework and analysis scripts, is available at https://github.com/AmelieNgo-research/Rmutaphy. Simulated datasets can be fully reproduced using the provided scripts. Hepatitis C virus data are publicly available from Smith et al. (2021).

## Acknowledgements

We thank our collaborators and colleagues at the MIVEGEC and LIRMM labs for helpful discussions and feedback. We also acknowledge the clinical teams involved in the hepatitis C cohort for data collection and management.

