## Supplementary material for "MutaPhy: A clade-based framework to detect genotype-phenotype associations on phylogenetic trees": SI

#### Supplementary Method S1 — Modeling background phenotype noise

In the main model, the probability of observing phenotype 1 is defined as

$$P(Z_s = 1 \mid IS_s) = \frac{1}{1 + \exp\{-k(IS_s - c) - r\}},$$

where $c$ and $k$ control the inflection point and the slope of the logistic function, respectively. Without additional constraints, this formulation implies a non-zero baseline probability when $IS_s = 0$.

To explicitly control this baseline probability, we introduce a parameter $b \in (0, 1)$ defined as

$$P(Z_s = 1 \mid IS_s = 0) = b.$$

Imposing this constraint yields

$$b = \frac{1}{1 + \exp\{-k(0 - c) - r\}} = \frac{1}{1 + \exp\{kc - r\}}.$$

Solving for $r$ gives:

$$\begin{aligned} & \frac{1}{1 + e^{-(-kc)-r}} = b \\ \iff \quad & 1 + e^{kc-r} = \frac{1}{b} \\ \iff \quad & e^{kc-r} = \frac{1}{b} - 1 \\ \iff \quad & r = kc - \log\left(\frac{1}{b} - 1\right) \end{aligned}$$

We considered two baseline-noise level in the simulations: a low-noise level with $b = 0.01$, corresponding to an almost negligible baseline probability, and a noisy level with $b = 0.2$, allowing for non-genetic background variability.

#### Supplementary Figure S1 — Effect of the proportion of causal mutations and background noise

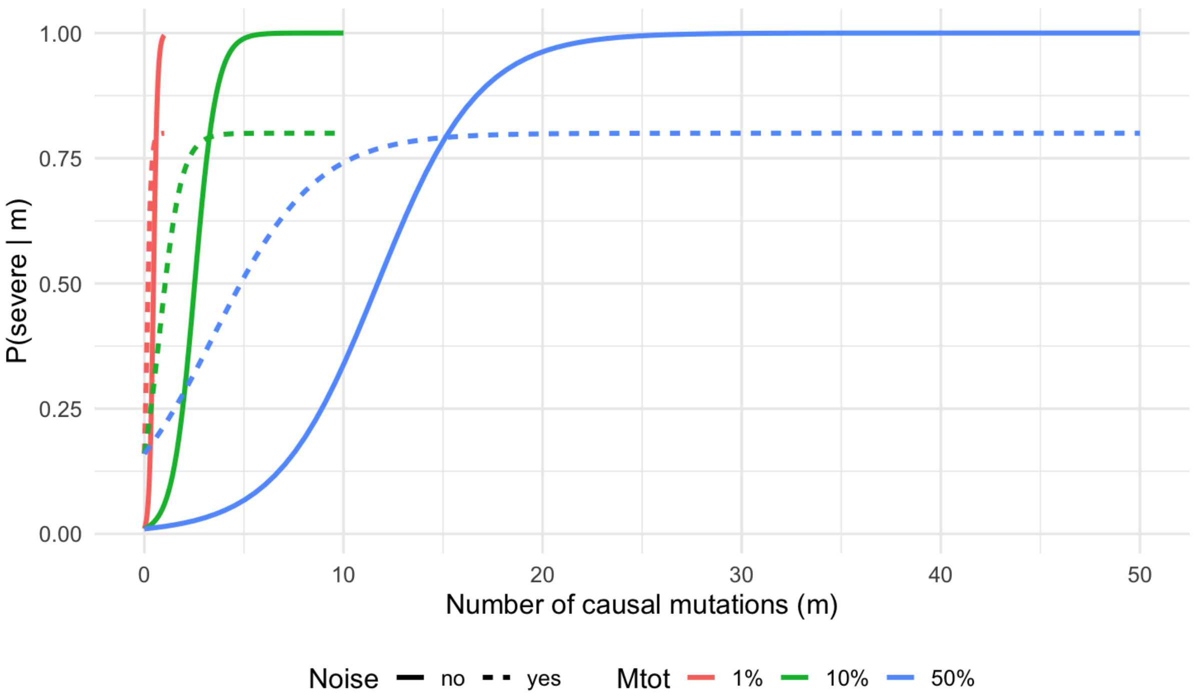


Figure 1: Effect of the causal mutation proportion $M_{\mathrm{tot}}$ and background noise on the probability of observing phenotype 1. The total number of simulated mutations is fixed to $N_{\mathrm{mut}} = 100$.

#### Supplementary Figure S2 — Subtree-level localization using raw permutation p-values

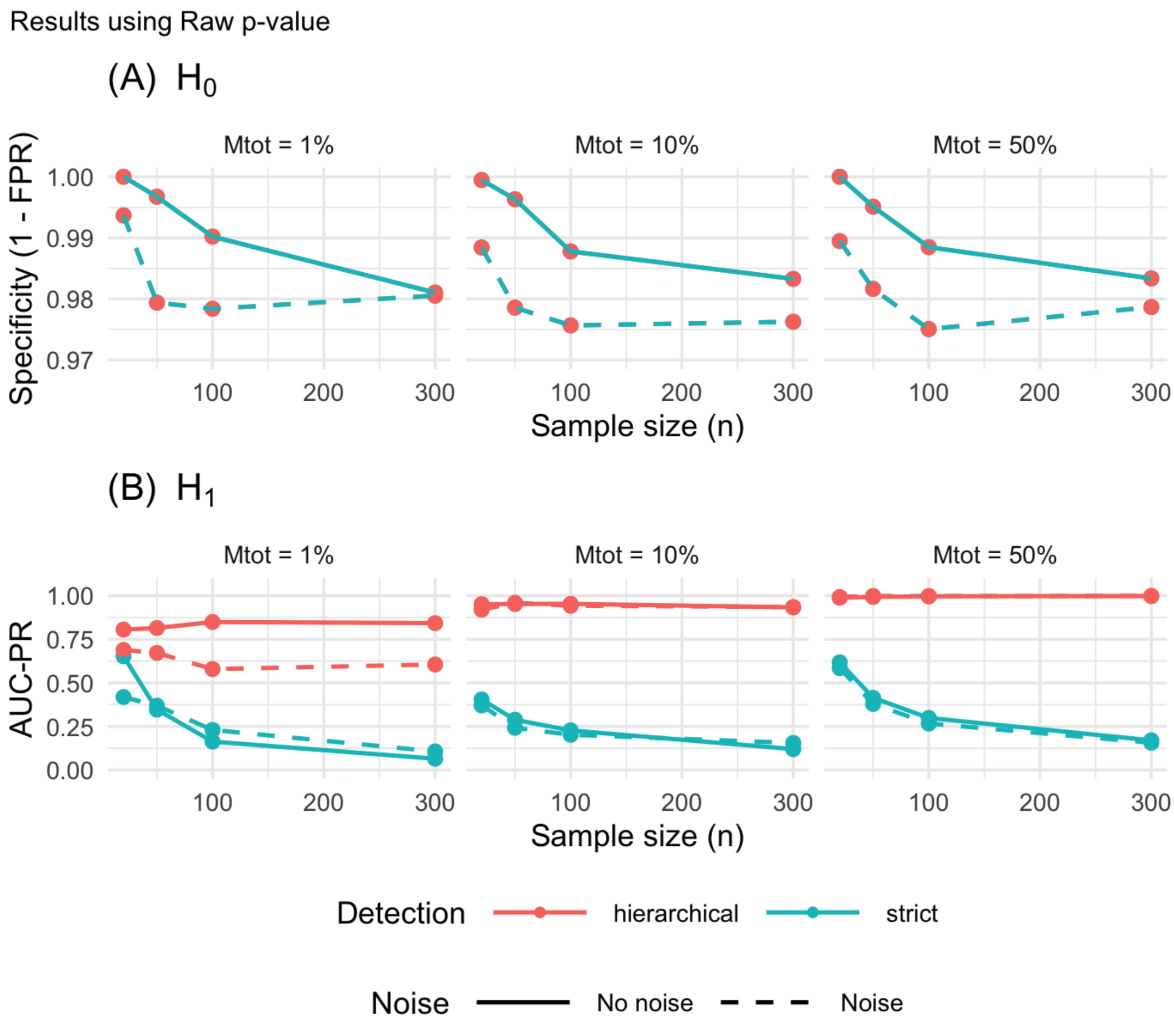


Figure 2: Subtree-level localization performance of MutaPhy using uncorrected permutation $p$-values, under strict and hierarchical evaluation definitions.

#### Supplementary Figure S3 — Comparison between permutation and hypergeometric subtree tests

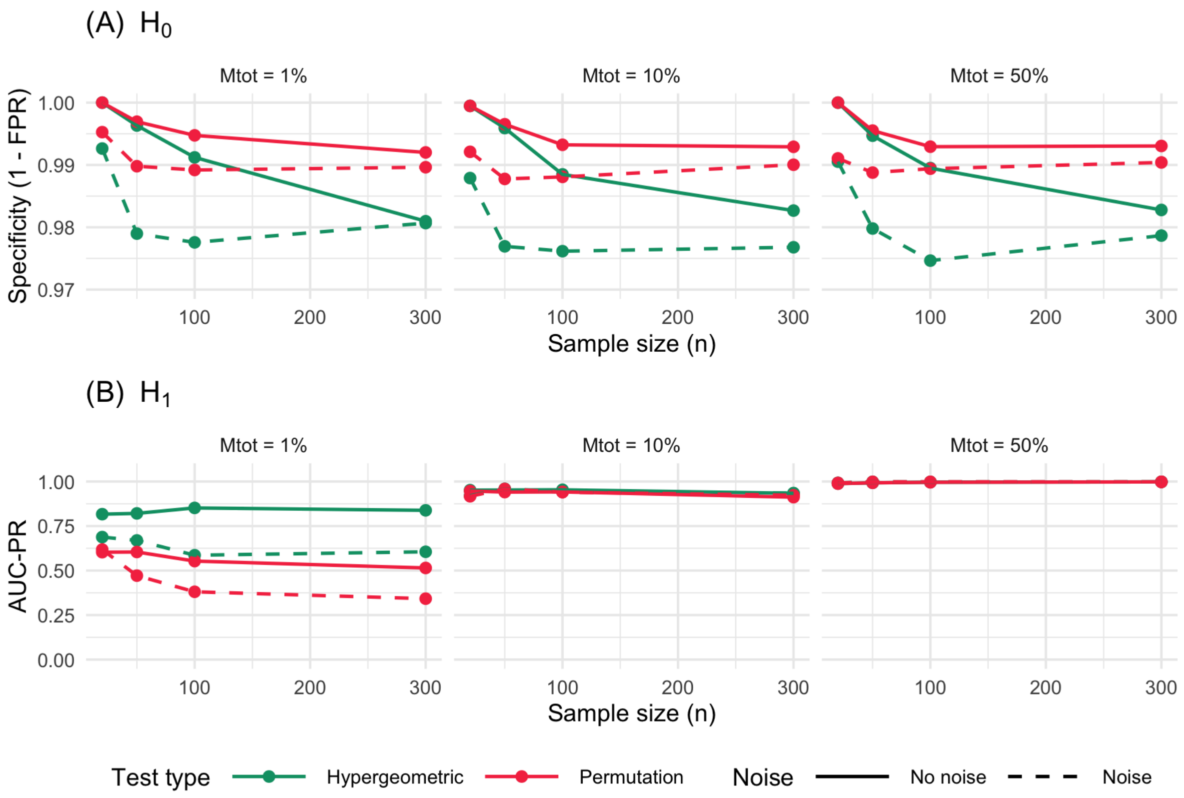


Figure 3: Comparison of subtree-level localization performance between the multiple-testing–corrected permutation test and the hypergeometric test under the hierarchical evaluation convention.

#### Supplementary Figure S4 — Precision of site-level candidate mutation detection by MutaPhy

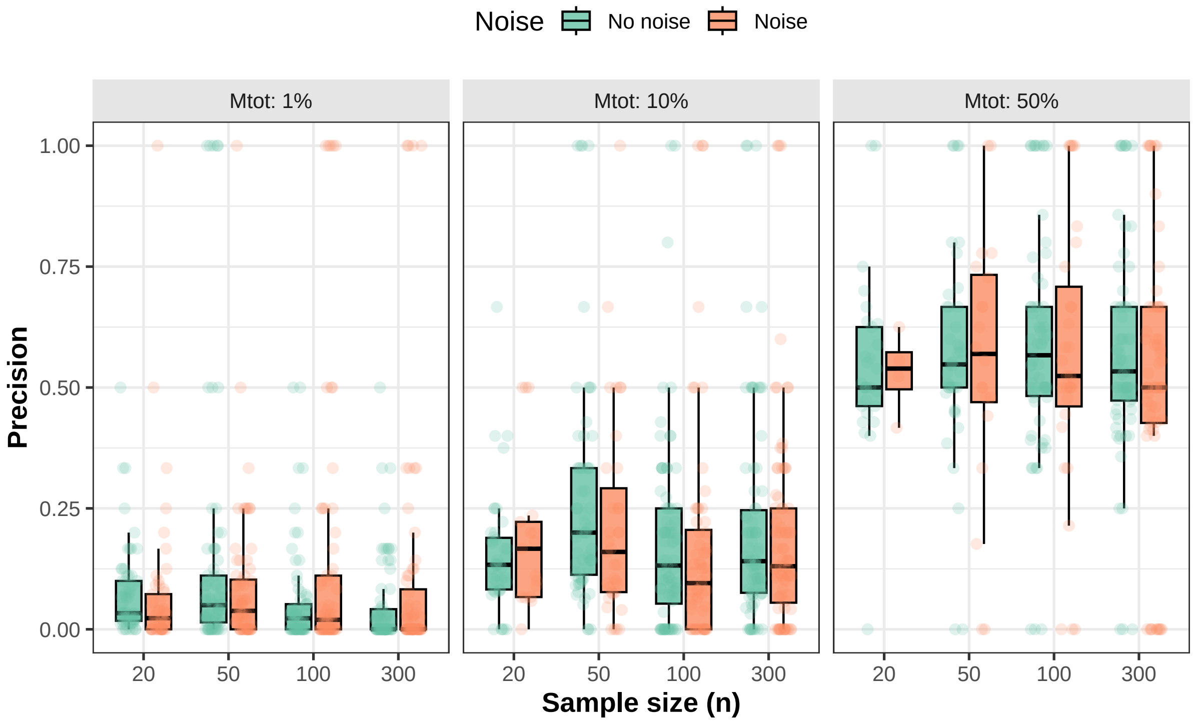


Figure 4: Precision of site-level candidate mutation detection by MutaPhy across simulation scenarios.

#### Supplementary Figure S5 — Recall of site-level candidate mutation detection by MutaPhy

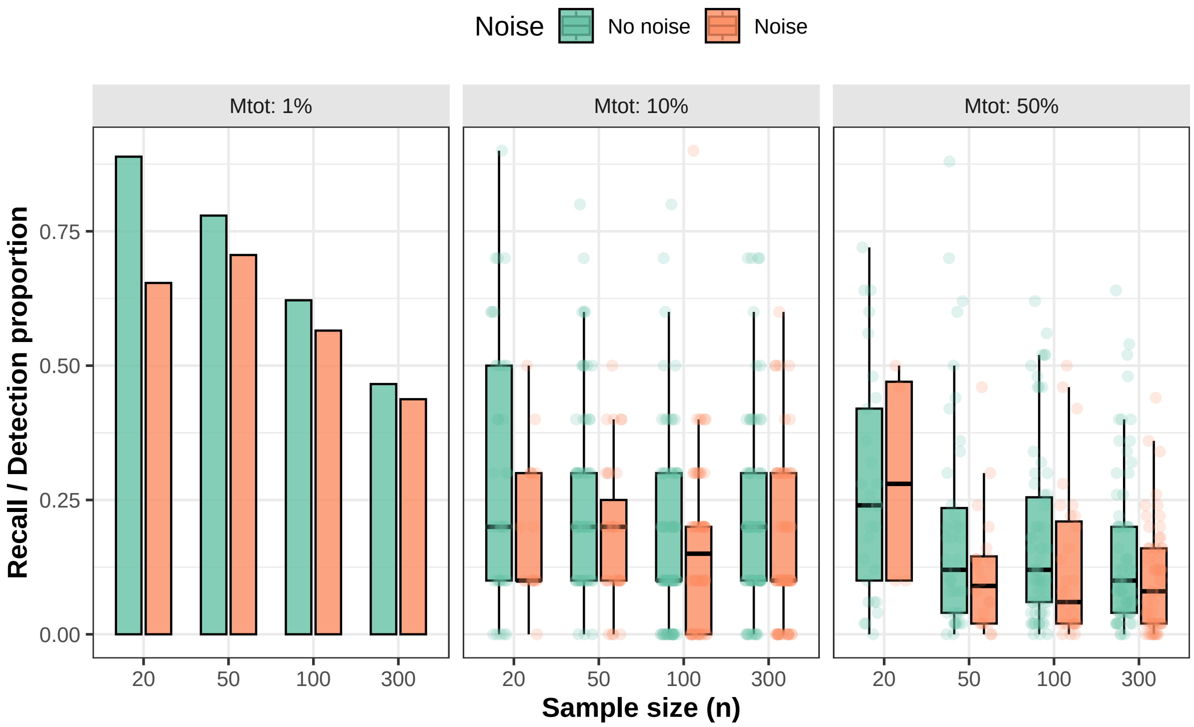


Figure 5: Recall of causal mutations among site-level candidates detected by MutaPhy.

### Supplementary Table 1 — Full simulation parameters

Table 1: Summary of simulation parameters

| Parameter | Values | Description |
|---|---|---|
| $N_{\mathrm{mut}}$ | 100 | Total number of simulated mutations per tree |
| $M_{\mathrm{tot}}$ | 1 %, 10 %, 50 % | Proportion of causal mutations among all simulated mutations |
| $n$ | 20, 50, 100, 300 | Number of tips (tree size) |
| Number of trees per scenario | 100 | Number of replicate trees simulated for each experimental condition |
| $\mu$ | $10^{-4}$ | Mutation rate per site and replication cycle |
| Branch scaling factor | 400 | Scaling factor applied to branch lengths to achieve 6–8% average sequence divergence |
| $\alpha$, $\beta$ | $\alpha = 4$, $\beta = 1$ | K80 substitution model parameters (transition and transversion rates) |
| $c$ | $\frac{M_{\mathrm{impact}}}{2}$ | Logistic midpoint (50% probability threshold for severe phenotype) |
| $k$ | $\frac{20}{M_{\mathrm{impact}}+1}$ | Logistic slope around the midpoint $c$ |
| $b$ | 0.01, 0.2 | Baseline probability of severe phenotype when $IS = 0$ (background noise level) |
| $r$ | Computed | Bias term enforcing $P(Z = 1 \mid IS = 0) = b$ |
| Mutation score ($H_0$) | 0 | Effect size assigned to mutations under the null hypothesis (no phenotypic effect) |
| Mutation score ($H_1$) | 1 | Effect size assigned to causal mutations under the alternative hypothesis |
| $N_{\mathrm{perm}}$ | 1000 | Number of permutations used to generate null distributions of association statistics |
| $\alpha$ (threshold) | 0.05 | Significance threshold for statistical tests |
